# Proprietary Processing of Placental Tissue Allografts Influences Wound Healing Trajectories in a Rodent Excisional Skin Wound Model

**DOI:** 10.64898/2026.09.11.748621

**Authors:** Ashley Robertson, Taylor Carson, Joydeep Basu

## Abstract

**Introduction:** Placental tissue allografts are widely utilized in the management of chronic wounds, including diabetic foot ulcers. In addition to serving as wound coverings, these products may influence wound repair through extracellular matrix architecture and biologically active signaling molecules native to human amniotic tissue. Manufacturing and processing methodologies vary considerably across commercially available products and may substantially impact biologic activity, tissue remodeling, and wound-healing trajectories. In this study, a rodent wound-healing model was utilized to compare a proprietary processed placental tissue allograft with a conventionally processed placental tissue comparator.

**Methods:** Two arms of a five-arm pilot rodent wound-healing study were evaluated to compare a proprietary processed human amniotic tissue allograft against a conventionally processed comparator; the remaining arms evaluated test articles under development and will be reported separately. Male rats (n=8 per arm), matched for weight, age, and diet, each received two full-thickness dorsal excisional wounds approximately 10 mm in diameter and 2 mm deep. Test articles were reapplied upon resorption over a 14-day period. At Day 14, sections were scored semi-quantitatively (0–4 Severity Injury Score) across histologic and immunohistochemical parameters spanning epidermal repair, inflammation, angiogenesis, proliferation, and remodeling. Cranial and caudal wounds were averaged within each animal, retaining the animal as the experimental unit. A Day 14 Histology Resolution Score was derived as a signed composite of pro- and anti-regenerative markers and evaluated alongside percent wound closure. Inferential statistics (Welch ANOVA with Games–Howell post hoc) were derived from the full five-arm cohort (n=40).

**Results:** Animals treated with the proprietary processed allograft showed more advanced progression toward wound resolution than those receiving the conventionally processed comparator, across both surface and tissue-level endpoints. Mean animal-level wound closure at Day 14 was 95.2% with the proprietary processed allograft versus 89.9% with the conventional comparator. The integrated Day 14 Histology Resolution Score — a composite of epidermal repair, inflammation, angiogenesis, proliferation, and remodeling endpoints — was 8.94 versus 5.13, corresponding to a shift from the poor-resolution range into the partial-resolution range on the pre-defined interpretive scale. Welch ANOVA across the full five-arm cohort (n=40) detected a significant overall effect of treatment on this composite (p = 0.010). The pairwise contrast between these two arms did not reach significance under Games-Howell correction across ten comparisons (p = 0.337); at n=8 per arm the study was not powered to resolve individual pairwise differences, and the observed standardized difference was large (Hedges’ g = 0.93). Injury endpoints were consistently lower in the proprietary processed allograft: 69% lower acute inflammation, 60% lower dermal necrosis, approximately 57% lower epithelial erosion, 46% lower ulceration, and 42% less hemorrhage. Importantly, these reductions in residual tissue injury occurred while maintaining robust granulation tissue formation, angiogenesis, collagen remodeling, cellular proliferation, and macrophage recruitment.

**Discussion:** These findings suggest that processing methodologies can materially influence the biologic performance of placental tissue allografts. The proprietary processed allograft was associated with lower residual tissue injury and greater regenerative and reparative activity relative to a conventionally processed comparator. These observations support the concept that placental tissue allografts should not necessarily be viewed as biologically equivalent and may have important implications for clinical outcomes and product selection in wound care.

## Introduction

Cutaneous wound healing is a tightly regulated biological process involving the coordinated interaction of inflammatory cells, resident stromal cells, extracellular matrix (ECM), soluble growth factors, and mechanical signals. Classically, wound repair progresses through four overlapping phases: hemostasis, inflammation, proliferation, and remodeling. Immediately following tissue injury, platelet activation initiates clot formation while simultaneously releasing cytokines and growth factors that recruit inflammatory leukocytes to the wound bed. During the inflammatory phase, neutrophils and monocyte-derived macrophages remove microorganisms and cellular debris while producing mediators that regulate the transition toward tissue repair. Successful resolution of inflammation is followed by the proliferative phase, characterized by fibroblast migration and proliferation, extracellular matrix deposition, angiogenesis, granulation tissue formation, and re-epithelialization mediated by keratinocyte migration. Finally, during remodeling, collagen fibers undergo maturation and reorganization, resulting in progressive restoration of tissue tensile strength while excessive matrix deposition is limited through balanced matrix metalloproteinase (MMP) and tissue inhibitor of metalloproteinase (TIMP) activity [1–5].

Chronic wounds fail to progress through this orderly sequence and instead become arrested in a prolonged inflammatory state. Persistent inflammation is associated with elevated concentrations of pro-inflammatory cytokines, excessive protease activity, degradation of endogenous growth factors, impaired angiogenesis, cellular senescence, and disruption of extracellular matrix architecture. The chronic wound microenvironment therefore lacks many of the biological cues necessary to support effective tissue regeneration, providing the rationale for advanced biologic therapies that restore a more favorable reparative environment [3–6].

Human amniotic membrane has been used clinically for more than one century. The earliest documented clinical application was described by Davis in 1910 for skin transplantation, while subsequent reports expanded its use for treatment of burns, chronic ulcers, ocular surface reconstruction, and surgical wound repair [7,8]. Interest in amniotic membrane has increased substantially over the past three decades following improved donor screening, tissue banking standards, and preservation technologies that permit safe manufacture of commercially available allografts. The biological properties of human amniotic membrane derive from its unique extracellular matrix composition and its reservoir of bioactive molecules. Native amnion contains multiple collagen isoforms (predominantly collagen types I, III, IV, V, and VII), laminin, fibronectin, hyaluronic acid, proteoglycans, and elastin, together with numerous cytokines and growth factors including epidermal growth factor (EGF), keratinocyte growth factor (KGF), hepatocyte growth factor (HGF), fibroblast growth factors (FGFs), platelet-derived growth factor (PDGF), transforming growth factor-β (TGF-β), and vascular endothelial growth factor (VEGF). Although the abundance of individual proteins varies considerably among donors and processing methods, collectively these extracellular matrix and signaling molecules provide structural support and biological cues that facilitate cell adhesion, migration, proliferation, angiogenesis, and matrix remodeling [9–13].

Current evidence suggests that amniotic membrane allografts accelerate wound repair through multiple complementary mechanisms rather than through delivery of a single active factor. First, the preserved extracellular matrix provides a provisional scaffold that supports fibroblast and keratinocyte attachment and migration. Second, matrix-bound cytokines and growth factors contribute to cellular proliferation and angiogenic responses. Third, amniotic membrane modulates inflammation by reducing excessive leukocyte activation and altering macrophage polarization toward a reparative phenotype in experimental models. Fourth, endogenous inhibitors of matrix metalloproteinases, including TIMP family proteins, together with suppression of excessive protease activity, may help preserve extracellular matrix integrity within chronic wounds. Finally, the membrane provides a protective biological covering that reduces evaporative fluid loss and serves as a temporary barrier against external mechanical injury while tissue regeneration proceeds [10–16].

Because native human amnion is mechanically fragile and biologically labile, numerous preservation and processing methods have been developed to facilitate commercial manufacture and clinical handling. Cryopreservation preserves viable or non-viable tissue architecture by minimizing ice crystal formation during controlled freezing, whereas dehydration methods—including lyophilization and low-temperature dehydration—permit ambient storage while maintaining much of the extracellular matrix structure. Additional manufacturing approaches include decellularization, gamma irradiation or electron-beam terminal sterilization, supercritical carbon dioxide processing, and chemical crosslinking intended to improve mechanical stability and resistance to enzymatic degradation [17–22]. Among these approaches, chemical modification has received particular attention because it can substantially alter the physical and biological characteristics of the extracellular matrix. Crosslinking agents such as glutaraldehyde, carbodiimide (EDC/NHS), genipin, and naturally occurring polyphenols including proanthocyanidins have been investigated to improve collagen stability, reduce degradation, and increase tensile strength. These reactions generally stabilize collagen by forming covalent intermolecular crosslinks between lysine or hydroxylysine residues within collagen fibrils. Although enhanced mechanical durability may prolong graft persistence within protease-rich chronic wound environments, excessive crosslinking may also reduce matrix porosity, alter degradation kinetics, decrease release of matrix-associated growth factors, modify cellular attachment sites, and impair host cell infiltration or remodeling. Consequently, the extent and chemistry of matrix modification represent important determinants of biological performance, requiring careful optimization to balance structural stability with preservation of regenerative bioactivity [20–27]. Given the diversity of commercially available processing technologies, relatively few studies have systematically compared how distinct manufacturing methodologies influence biological performance in vivo. The present study evaluates a proprietary human amnion processing methodology relative to conventionally processed human amnion allografts using a standardized rodent full-thickness excisional wound model. By comparing wound closure kinetics together with histological and tissue remodeling endpoints, this work seeks to determine whether differences in tissue processing translate into measurable differences in regenerative performance following implantation.

## Materials & Methods

Rodent studies were performed at Bio-Legacy Research, San Diego. *n*=8 male Sprague-Dawley rats matched for age, weight, diet and wound size were assigned to each of five experimental test groups. This report presents two: (A) conventionally processed human amniotic tissue allograft and (B) proprietary processed human amniotic tissue allograft, XWRAP. The remaining three arms evaluated test articles under active development and will be reported separately. The Group A versus Group B comparison addressed a pre-specified hypothesis of the original five-arm design concerning the effect of processing methodology on wound healing trajectory.

Each rodent underwent two full thickness skin excisional wounds on the dorsal surface, with one wound on the cranial side and one wound on the caudal side [28]. Each wound was approximately 10mm in diameter and 2mm deep. Both wounds on each animal were covered with the treatment allograft and further secured with gauze and bandages to hold the test article in place. Rodents were jacketed and additionally wrapped to minimize disturbance of the wound during the study period. Wounds were inspected and documented every 48 hours, with the integrity of the test article being monitored throughout the healing process. Upon complete resorption of a test article, a new test article was implanted into the wound bed to accurately mimic current clinical recommendations for the use of XWRAP in treatment of chronic wounds. Rodents were kept on study for 14 days and euthanized humanely.

Upon necropsy, a representative sample was excised from each wound (cranial and caudal sites). Once the samples were excised, the cranial section was inked for histological identification, before being placed in a standard tissue cassette for fixation in 10% neutral buffered formalin (NBF) for no less than 48-72 hours, followed by paraffin processing and staining per Bio-legacy’s internal SOPs. The remaining skin (wound) specimen was preserved in 10% NBF in a properly labeled container. Trimmed sections were submitted for paraffin processing and embedding. Paraffin blocks were cut to a 4-5μm thickness using a rotary microtome. Sections were mounted on glass slides and stained with Hematoxylin and Eosin (H&E) and Masson’s trichrome, Picro-Sirius Red [29] and five additional immunohistochemistry stains selected to characterize key mechanistic axes of the wound healing process: CD31, Ki-67, CD45, CD68 and CD163 [30,31,32,33].

### Histological Evaluation

Stained slides were transferred to Infinium Pathology Consultants, 8215 Messenger Ct. Stokesdale, NC 27357 for microscopic evaluation. MS Excel was utilized for data capture and basic analysis (Mean ± SD) and submitted to Applied Biologics for further statistical analysis. Histological assessment was performed by light microscopy. All sections were semi-quantitatively scored by a single observer unblinded to the treatment group using the following parameters: Re-epithelialization (epidermal coverage), epidermal thickening (acanthosis), papillary dermal edema, papillary dermal necrosis, granulation tissue, acute inflammation, giant cell reaction, erosion, ulceration and hemorrhage. These parameters were semi-quantitatively scored by assigning a severity injury score (SIS) graded from “0” through “4” based upon the criteria listed in Table 1 below.

**Table 1.** Severity Injury Score (SIS): grading criteria used for histopathology review.

| Grade | Interpretation |
| --- | --- |
| 0, Not present | No meaningful histologic change from normal, or not scored |
| 1, Minimal | Minor or infrequent change, involving less than 10 percent of the tissue |
| 2, Mild | Clearly noticeable but not prominent change, involving about 10 to 25 percent of the tissue |
| 3, Moderate | Prominent and consistently present change, involving about 26 to 50 percent of the tissue |
| 4, Marked | Overwhelming or persistent change, involving more than 50 percent of the tissue |

In addition, the five immunohistochemistry-stained sections were evaluated/scored. The assessment and grading of immunohistochemistry (IHC) stained sections for the expression markers (CD31, Ki-67, CD45, CD68 and CD163), was conducted unblinded by a single observer. All slides were examined at low and high-power magnification for adequacy of tissue elements and staining. The intensity of staining was graded as follows: 0 (negative), 1+ (weak), 2+ (mild), 3+ (moderate-strong), and 4+ (intensely strong).

### Surface Closure Analysis (Dataset Preparation and Methodology)

The wound closure dataset was structured as repeated wound-area measurements for each animal, wound site, treatment group, and study day. Data cleaning was performed to standardize identifiers, preserve wound-site labels, and reshape the dataset into a format suitable for plotting and per-animal summaries. For each wound, Day 0 area was used as the baseline reference. Normalized wound area was calculated by dividing each later wound-area measurement by the corresponding Day 0 value. Percent closure was then calculated as:

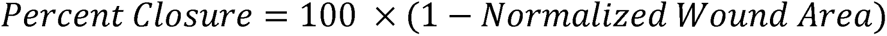

This approach allowed wounds with different baseline sizes to be compared on the same relative scale. Both wound-site-level and animal-level summaries were prepared. Cranial and caudal wounds were retained separately for site-comparison analyses. Prior to defining the primary closure summary, cranial and caudal wounds were compared directly to determine whether wound site should be retained as an analytical factor. Paired site-level comparisons showed some animal-level variation in both direction and magnitude but no consistent directional site bias across treatment arms. On this basis, a MeanSites summary was generated by averaging the paired cranial and caudal values within each animal. This preserved the animal as the experimental unit while reducing site-level noise in the absence of a stable site effect. Time-window summaries were derived to capture cumulative healing over the full study window, days 0-14.

### Histopathology Analysis (Histology Dataset and SIS Scoring Framework)

The histopathology dataset contained semi-quantitative tissue scores at day 14, time of necropsy. Tissue sections were reviewed using a Severity Injury Score (SIS) scale from 0 to 4, where larger values reflected broader or more persistent tissue involvement. This framework allowed different tissue features to be placed on a common semi-quantitative scale while preserving biological context.

### Integrated Surface Closure and Histopathology Analysis

Surface wound closure is an important and intuitive measure of healing, but it does not fully describe the biological state of the tissue beneath the surface. A wound may appear largely or completely closed while still differing in inflammation, remodeling, collagen organization, residual injury, or retained material. For this reason, day 14 wound closure was integrated with matched day 14 histopathology to evaluate whether surface-level healing aligned with tissue-level resolution. The central goal of this analysis was to treat wound closure as a measurable proxy for healing rather than assuming it fully represents biological repair. Longitudinal wound-closure data and terminal histopathology were available from the same animals, which made it possible to directly compare macroscopic closure with microscopic tissue status. This integration was a key focus of the analysis because it allowed treatment performance to be evaluated from both the visible wound surface and the underlying histological response.

The two datasets were merged on animal identifier at the day 14 timepoint, pairing each animal’s MeanSites closure value with its day 14 histopathology, producing an animal-level dataset containing both endpoints. To support interpretation, histology markers were grouped into three categories based on their interpretation at the terminal timepoint: pro-regenerative markers, anti-regenerative markers, and mixed/context-dependent markers. This grouping was used because the biological meaning of a given feature depended not only on its magnitude but also on whether it reflected repair, unresolved injury, or a stage-dependent process. A signed SIS framework was constructed so that markers associated with more regenerative tissue behavior contributed in a positive direction and markers associated with more adverse or unresolved tissue behavior contributed in a negative direction (Table 2). Mixed markers were intentionally withheld from the signed score so that their meaning would not be oversimplified. A key mixed feature was the M1/M2 macrophage ratio, calculated as CD68 divided by CD163 [33]. This ratio was used as an indicator of macrophage polarization and wound phase.

**Table 2:**
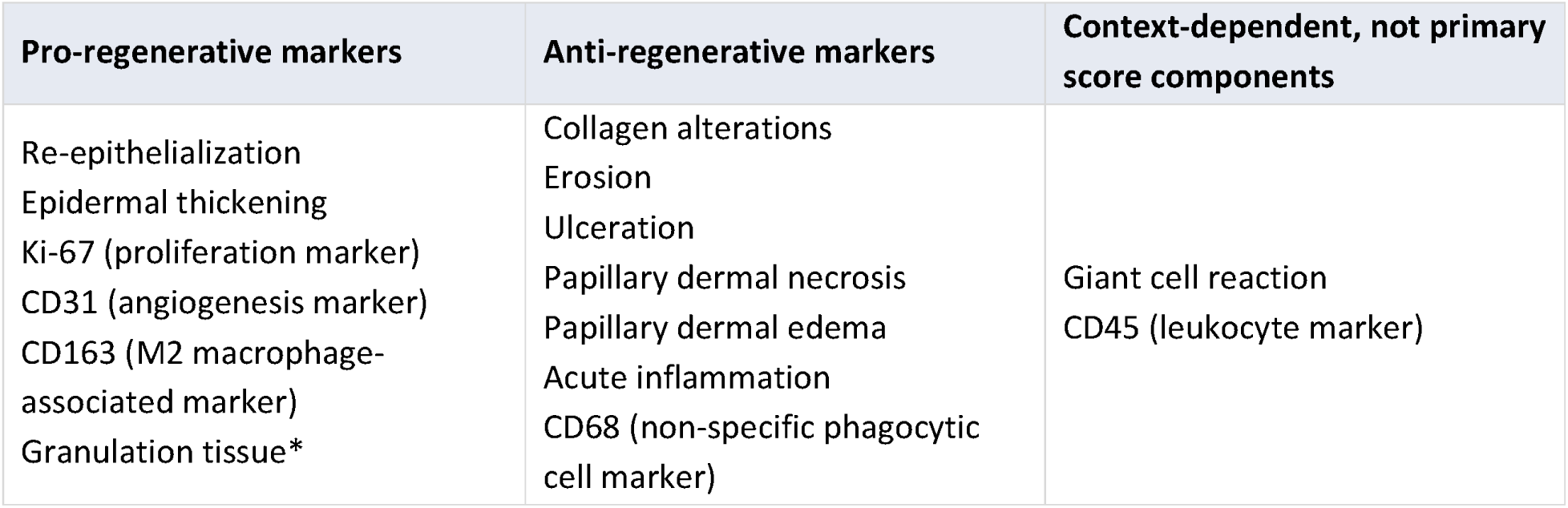
Day 14 histology marker interpretation used for the integrated resolution score.

| Pro-regenerative markers | Anti-regenerative markers | Context-dependent, not primary score components |
| --- | --- | --- |
| Re-epithelialization<br>Epidermal thickening<br>Ki-67 (proliferation marker)<br>CD31 (angiogenesis marker)<br>CD163 (M2 macrophage-associated marker)<br>Granulation tissue* | Collagen alterations<br>Erosion<br>Ulceration<br>Papillary dermal necrosis<br>Papillary dermal edema<br>Acute inflammation<br>CD68 (non-specific phagocytic cell marker) | Giant cell reaction<br>CD45 (leukocyte marker) |

Granulation tissue was treated as pro-regenerative, but high Day 14 granulation was not interpreted as the most resolved remodeling state. Therefore, a modified contribution was used for this marker.

The composite histology resolution score was calculated for each animal as:

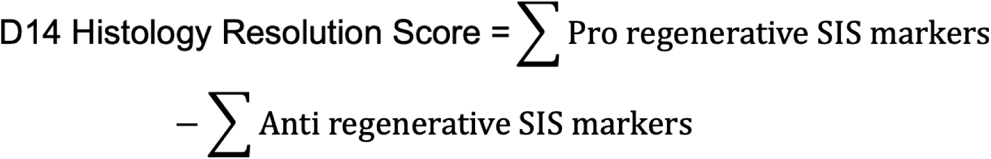

All markers contributed with equal weight and without rescaling. In consultation with the evaluating pathologist, both absent granulation (SIS 0) and persistently high granulation (SIS 4) were assessed as unfavorable states at day 14 relative to a resolving wound, the former indicating failed repair and the latter ongoing rather than completed repair. Its contribution was therefore calculated as:

Higher day 14 Histology Resolution Scores indicate a more favorable tissue-level healing profile. Lower values indicate greater evidence of unresolved injury, inflammation, or incomplete remodeling. The observed day 14 score range in this dataset ranged from −1.0 to 14.5. By construction, values above 0 indicate that pro-resolution features outweighed unresolved-injury features (Table 3).

**Table 3:**
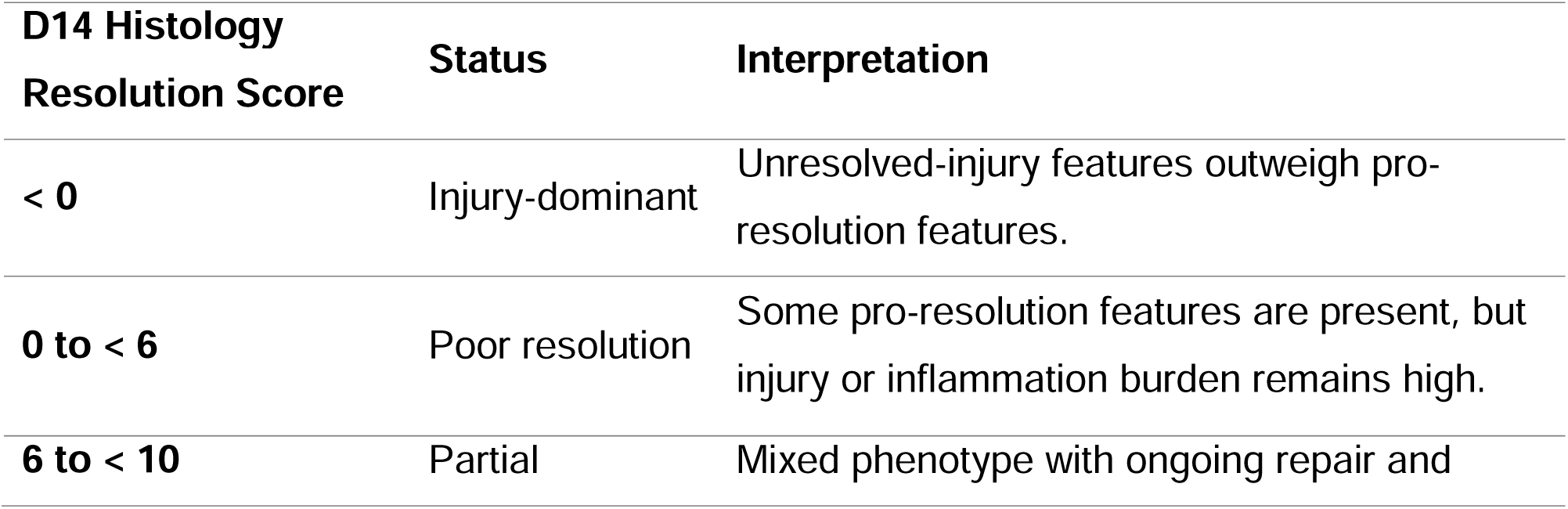
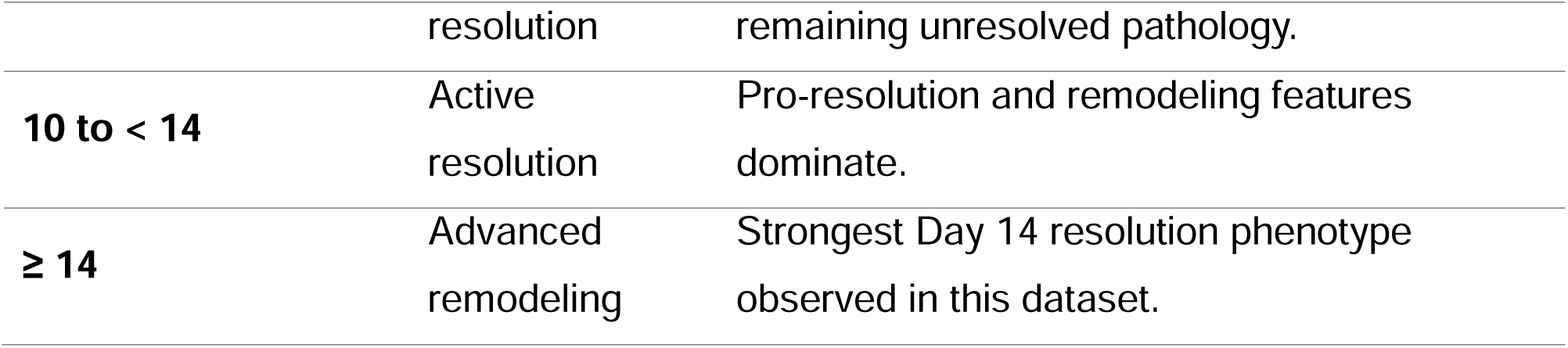
Interpretive labels used for the day 14 Histology Resolution Score.

## Statistical Analysis

Analyses were performed in Python 3.12.3 using pandas 2.2.2, NumPy 1.26.4, SciPy 1.13.1, and pingouin 0.5.5. Given small group sizes and unequal variance across treatment arms, the overall treatment comparison used Welch’s ANOVA, followed by Games-Howell post hoc pairwise contrasts. These tests were applied to the day 14 Histology Resolution Score across all five treatment arms (N=40). The omnibus test indicated a significant overall effect of treatment (*F*(4, 17.2) = 4.67, *p*=0.010, 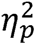 = 0.29). The Group A versus Group B contrast is reported here, and the remaining contrasts involve arms presented separately. All inferential statistics reported in this manuscript were derived from the complete five-arm day 14 cohort (N=40) and are not recalculable from the two arms presented. The association between day 14 percent closure and Histology Resolution Score was assessed by Pearson correlation across the full five-arm cohort (n=40).

Individual histology markers were compared between Groups A and B descriptively (mean ± SD and percent difference) and were not tested inferentially; no marker-level p-values are reported. Group-level z-scores were calculated across the five treatment-arm means using SciPy zscore. They express each arm’s position relative to the dispersion of five arm means.

## Results

Representative histological and immunohistochemical sections from Group A and Group B at Day 14 further illustrate the treatment-level differences summarized above (Figure 1, Group A; Figure 2, Group B). Low-power H&E and Masson’s trichrome sections showed a more advanced wound-closure phenotype in Group B, with complete epithelial coverage and only scant residual scab (Figure 2A–B), compared with near-complete but less mature epithelial coverage overlying a more prominent scab in Group A (Figure 1A–B), consistent with the higher re-epithelialization score observed in Group B (3.75 ± 0.45 vs. 3.44 ± 0.63). Masson’s trichrome and Picro-Sirius Red staining under polarized light showed a more mature, Type I–predominant collagen organization in Group B relative to Group A (Figures 1C, E and 2C, E), a visual pattern that accompanied closely matched semi-quantitative collagen alteration scores between groups (2.63 ± 0.50 vs. 2.69 ± 0.48). Ki-67 immunolabeling confirmed active, focal proliferation at the dermo-epidermal junction in both groups, with a modestly higher proliferative index in Group B (1.69 ± 0.60 vs. 1.50 ± 0.63; Figures 1D, 2D). CD31 staining revealed moderate-to-marked micro-vessel density in both groups (Figures 1F, 2F), indicating that angiogenic support of the healing wound bed was preserved regardless of processing method.

**Figure 1:**
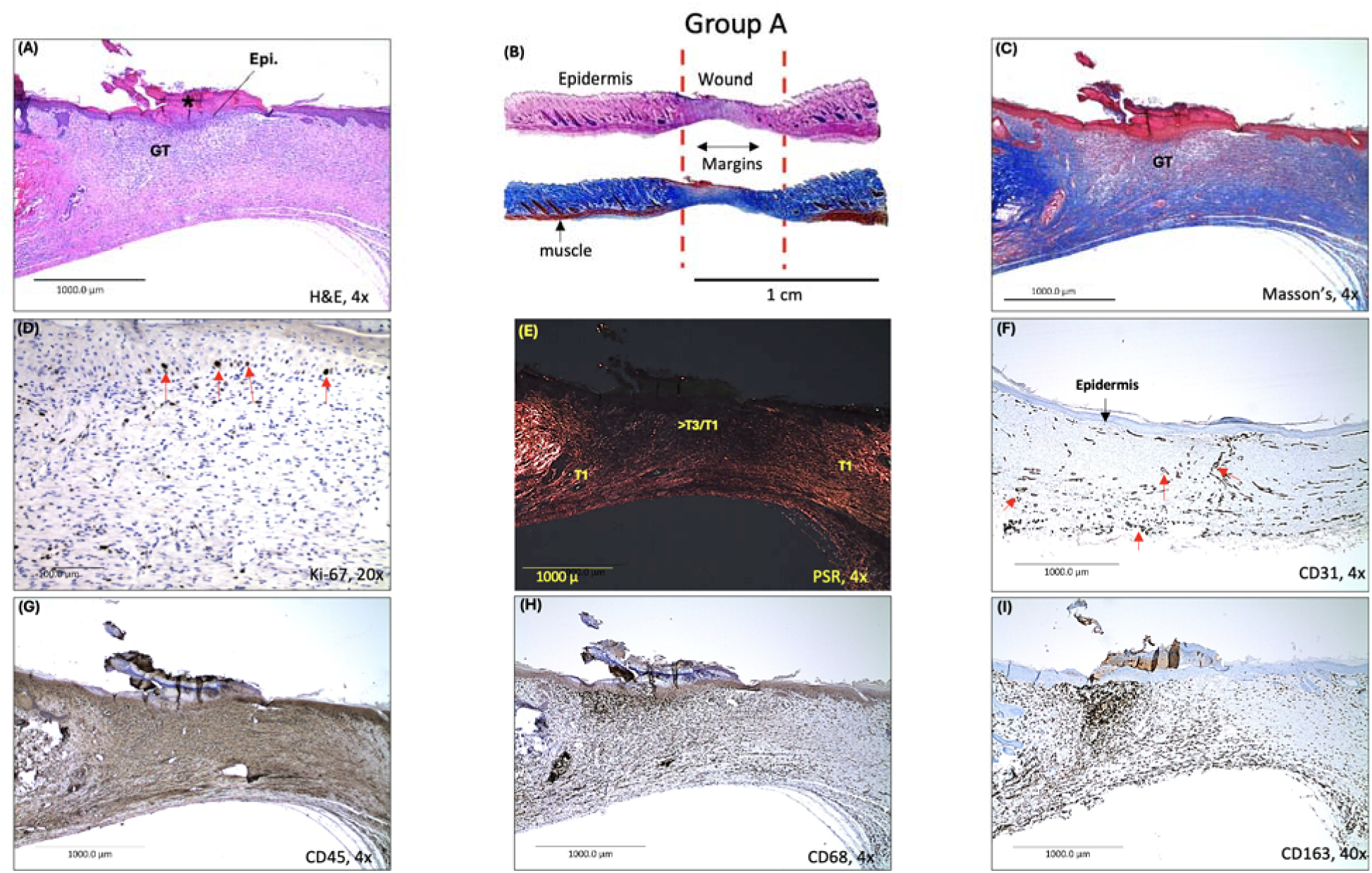
Representative histological and immunohistochemical evaluation of wound healing in Group A. **(A)** Low-power H&E section showing near-complete epithelial (Epi.) coverage over the defect with overlying scab formation (*) and underlying moderate granulation tissue (GT) in the upper dermis. **(B)** Low-power magnification (H&E and Masson’s trichrome) highlighting the wound and defect margins (dashed lines). *Note: Scale bar applies only to this panel as indicated.* **(C)** Masson’s trichrome section demonstrates remnant granulation tissue (GT) and mild-to-moderate collagen deposition (blue), with slightly less density than that observed in Group B. **(D)** Ki-67 immunohistochemistry (IHC) showing focal cell proliferation via dark-brown nuclear staining (arrows) at the dermo-epidermal junction. **(E)** Picro-Sirius Red (PSR) staining under polarized light, distinguishing mature Type I **(T1)** collagen (orange-red) from immature/thinner Type III **(T3)** fibers (greenish-yellow). **(F)** CD31 IHC highlighting endothelial cell membranes and cytoplasm (red arrows), evidence of moderate to marked, micro-vessel (angiogenesis) density. **(G–I**) Inflammatory cell infiltration is characterized by brown staining (membranous and cytoplasmic) expression: **(G)** CD45 (total white blood cells) exhibiting intensely strong expression, indicating a high presence of leukocytic infiltrate. (**H)** CD68 (M1/M2 macrophages), and **(I)** CD163 (M2 macrophages) both markers show moderate-strong expression, indicative of M2-dominated remodeling.

**Figure 2:**
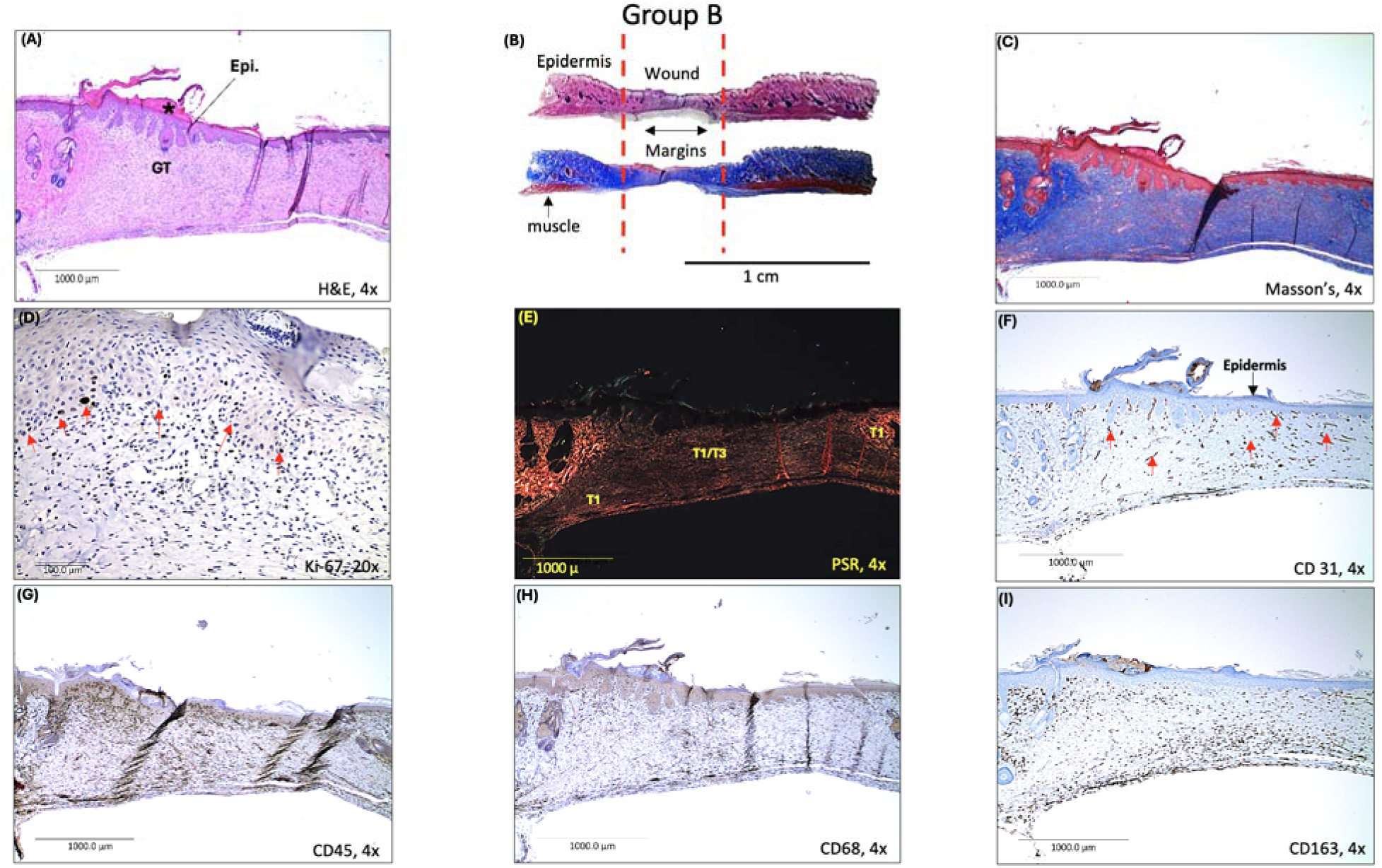
Representative histological and immunohistochemical evaluation of wound healing in Group B. **(A)** Low-power (4X) H&E section showing complete epithelial (Epi. coverage of the defect with scant overlying scab formation (*), an overall wound-surface closure/re-epithelialization score of 3.75 ± 0.45 versus 3.44 ± 0.63 in Group A, and mild to moderate granulation tissue (GT) in the dermis. **(B)** Low-power H&E and Masson’s trichrome images highlighting the wound and defect margins (dashed lines). *Note: The scale bar applies only to this panel, as indicated.* **(C)** Masson’s trichrome section demonstrating greater collagen deposition density (blue) than observed in Group A. (**D**) Ki-67 immunohistochemistry (IHC) showing focal cell proliferation, indicated by dark-brown nuclear staining (arrows), at the dermo-epidermal junction, with a slightly higher favorable mean ± SD score in Group B vs. Group A (1.69 ± 0.60 vs. 1.50 ± 0.63). **(E)** Picro-Sirius Red (PSR) staining under polarized light distinguishing mature Type I (T1) collagen (orange-red) from immature Type III (T3) fibers (greenish-yellow), with collagen alterations slightly higher in Group A vs. Group B (2.69 ± 0.48 vs. 2.62 ± 0.50). **(F)** CD31 IHC highlighting endothelial cell membranes and cytoplasm (red arrows), consistent with moderate to marked micro-vessel (angiogenesis) density. **(G–I)** Inflammatory cell infiltration showing mild to moderately strong brown membranous and cytoplasmic expression: **(G)** Total leukocytic infiltration, quantified via CD45 expression, remained similarly robust between Group A (3.56 ± 0.51) and Group B (3.63 ± 0.50) at Day 14, indicating a sustained, active immune cell presence in both treatment cohorts. However, the downstream functional outcomes diverged drastically; the persistent inflammatory burden in Group A correlated with localized tissue degradation, whereas Group B efficiently channeled this leukocyte pool toward tissue repair mechanisms. **(H)** CD68 (M1/M2 macrophages): The overall dermal macrophage population using CD68 pan-marker staining showed identical absolute recruitment levels in both groups (3.19 ± 0.83). While the physical abundance of macrophages at the injury site did not differ, Group B utilized this cellular framework to resolve active, destructive inflammation— significantly reducing acute surface inflammation scores by **69.3%** and dermal necrosis by **60.0%** relative to Group A. **(I)** CD163+ expression demonstrated robust M2 macrophage activation across both cohorts, with Group A (3.38 ± 0.62) and Group B (3.13 ± 0.62) maintaining high concentrations relative to the total pan-macrophage pool (CD68: (3.19). Importantly, while Group A exhibited a slightly higher raw density of CD163+ cells, this immune profile was counter-productive, coexisting with unresolved tissue degradation, elevated necrosis (1.25 ± 1.13) vs. Group B (0.50 ± 0.73) and extensive ulceration (1.63 ± 1.54) vs. Group B (0.88 ± 1.26). In contrast, Group B leveraged a more efficient, tightly regulated M2 response to quiet the local micro-environment, driving a 69% reduction in acute inflammation and a 60% reduction in necrosis, which ultimately facilitated definitive epidermal barrier restoration.

The immune cell infiltrate provided the clearest histological distinction between groups. Total leukocytic infiltration (CD45) and overall macrophage recruitment (CD68) were quantitatively similar between Group A and Group B (CD45: 3.56 ± 0.51 vs. 3.63 ± 0.50; CD68: 3.19 ± 0.83 in both groups; Figures 1G–H, 2G–H), indicating that both treatments elicited a comparably robust immune response. However, the functional consequence of this response diverged sharply between groups. In Group A, the persistent inflammatory infiltrate coexisted with unresolved tissue injury, reflected by higher rates of dermal necrosis (1.25 ± 1.13 vs. 0.50 ± 0.73) and ulceration (1.63 ± 1.54 vs. 0.88 ± 1.26) despite a slightly higher density of CD163+ M2 macrophages (3.38 ± 0.62 vs. 3.13 ± 0.62; Figure 1I). In Group B, by contrast, a comparable leukocyte and macrophage population was more efficiently channeled toward inflammatory resolution rather than ongoing tissue damage, evidenced by a 69.3% reduction in acute inflammation and a 60.0% reduction in dermal necrosis relative to Group A (Figure 2G–I). Taken together, these representative histological and immunohistochemical findings support the conclusion that the proprietary processing methodology (Group B) did not simply suppress the immune response, but rather promoted a more productive, pro-resolving inflammatory phenotype that translated into a more advanced Day 14 wound-healing trajectory relative to conventional processing (Group A). Viewed collectively, the Day 14 findings reveal a consistent treatment-level pattern. The proprietary processed allograft maintained the major regenerative processes required for wound repair while exhibiting substantially less residual tissue injury than the conventionally processed comparator. Granulation tissue formation was equivalent between groups, while angiogenesis, collagen remodeling, cellular proliferation, and macrophage recruitment remained robust. Against this background of preserved regenerative activity, the proprietary processed allograft demonstrated 69% lower acute inflammation, 60% lower dermal necrosis, approximately 46% lower ulceration, and approximately 57% lower epithelial erosion. Thus, the observed difference was not simply suppression of the wound-healing response, but rather preservation of regenerative activity in association with substantially reduced unresolved injury and inflammatory burden.

A summary of the overall histopathological outcomes (*n*=8) upon end of study is presented in Table 4 below.

**Table 4:** Day 14 histology marker outcomes by treatment group, presented as mean ± SD (n=8 animals per group).

Epidermal Findings
| Parameter | Group A<br>(Mean ± SD) | Group B<br>(Mean ± SD) | Notes |
| --- | --- | --- | --- |
| Re-epithelialization | 3.44 ± 0.63 | 3.75 ± 0.45 | Strong in both; Group B shows excellent resurfacing. |
| Epidermal thickening<br>(Acanthosis) | 3.00 ± 0.63 | 3.50 ± 1.03 | Group A: most pronounced hyperplasia, reflecting heightened epithelial turnover.<br>Group B: high epidermal activity; larger SD suggests uneven epithelial maturity across animals. |
| Erosion | 0.88 ± 0.50 | 0.38 ± 0.62 | Minimal to mild average severity in both groups. |
| Ulceration | 1.63 ± 1.54 | 0.88 ± 1.26 | Group A: variable severity (minimal–moderate) by animal and wound location; average minimal–mild. Group B: scattered superficial defects, minimal papillary dermis involvement. |

Dermal Findings
| Parameter | Group A<br>(Mean ± SD) | Group B<br>(Mean ± SD) | Notes |
| --- | --- | --- | --- |
| Edema | 1.00 ± 1.03 | 1.06 ± 1.12 | Minimal in both; Group A associated with mild acute inflammation. |
| Necrosis | 1.25 ± 1.13 | 0.50 ± 0.73 | Group A: minimal, likely linked to mild acute inflammation. Group B: variable (0–3 range) but overall minimal (1+). |
| Granulation tissue | 2.94 ± 0.25 | 2.94 ± 0.25 | Identical; moderately high in Group A, consistent with active normal wound progression. |
| Collagen alterations | 2.69 ± 0.48 | 2.63 ± 0.50 | Mild–moderate in both; active remodeling with appreciable type I collagen. |
| CD31 | 4.00 ± 0.00 | 3.94 ± 0.25 | Group A: abundant angiogenesis, uniformly high. Group B: vigorous angiogenesis, possibly driven by persistent tissue stress. |
| Ki-67 | 1.50 ± 0.63 | 1.69 ± 0.60 | Comparable between groups; consistent with ongoing epithelial proliferation. |
| Hemorrhage | 0.53 ± 0.63 | 0.31 ± 0.48 | Minimal in both; upper dermis, attributed to fragile capillaries in developing granulation tissue, ulceration, and/or dressing changes. |

Inflammation
| Parameter | Group A<br>(Mean ± SD) | Group B<br>(Mean ± SD) | Notes |
| --- | --- | --- | --- |
| Acute inflammation | 1.63 ± 1.15 | 0.50 ± 0.73 | Group A: mild, confined to papillary dermis. Group B: minimal, confined to wound surface (scab formation). |
| Foreign-body giant cell<br>response | Slightly elevated<br>(qualitative) | Slightly elevated<br>(qualitative) | Both attributed to incidental rat hair introduction during surgery or dressing/test article reapplication. |

Immunohistochemical Markers
| Parameter | Group A<br>(Mean ± SD) | Group B<br>(Mean ± SD) | Notes |
| --- | --- | --- | --- |
| CD45 | 3.56 ± 0.51 | 3.63 ± 0.50 | Strong leukocytic infiltrate in both. |
| CD68 | 3.19 ± 0.83 | 3.19 ± 0.83 | Identical. |
| CD163 | 3.38 ± 0.62 | 3.13 ± 0.62 | Group A: M2-dominated remodeling. Group B: moderate expression, comparable to most groups, indicating active matrix remodeling. |

The combined day 14 histology resolution score for groups A and B is summarized as Table 5 below. (update this to table 5 in source table)

**Table 5.** Group B (proprietary processing) outperforms Group A (standard processing) across all four metrics – higher histology resolution score, higher % closure, and both z-scores shift from negative (below cohort average) to positive (above average).

| Group | Treatment | D14 Histology Resolution Score | % Closure | Histology Z | Closure Z |
| --- | --- | --- | --- | --- | --- |
| A | Standard processing | 5.125 | 89.935 | -1.469 | -1.520 |
| B | Proprietary processing | 8.938 | 95.174 | 0.172 | 0.395 |

Group B showed a higher mean day 14 Histology Resolution Score than Group A (8.94 vs. 5.13, a difference of 3.81 points), corresponding to a shift from the poor resolution range into the partial resolution range on the interpretive scale (Table 3). Welch ANOVA across the full five-arm cohort detected a significant overall effect of treatment on the composite score (*F*(4, 17.2) = 4.67, *p* = 0.010, = 0.29). The Group A versus Group B pairwise contrast did not reach significance under Games-Howell correction across ten comparisons (p = 0.337), with a large standardized effect size (Hedges’ g = 0.93); this pilot was not powered to resolve individual pairwise differences. Animal-level scores are shown in Figure 3. Group A included multiple animals in the injury-dominant and poor-resolution ranges, while Group B animals were distributed across the partial- and active-resolution ranges.

**Figure 3:**
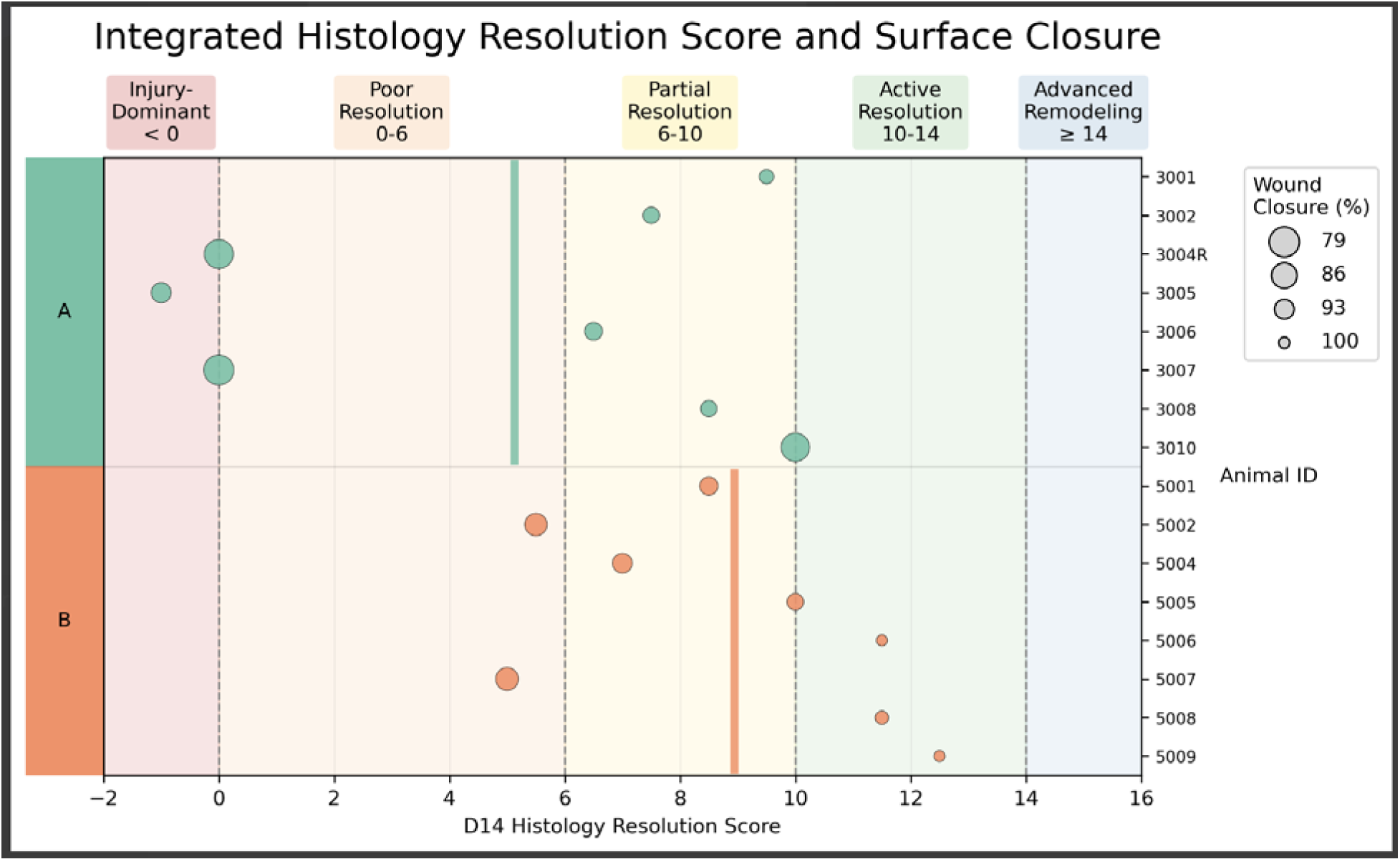
Day 14 Histology Resolution Score by animal, Groups A, B. Each point represents one animal. The x-axis shows the day 14 Histology Resolution Score, with shaded regions indicating injury-dominant, poor resolution, partial resolution, active resolution, and advanced remodeling ranges. Vertical colored bars indicate treatment-level mean resolution score. Bubble size represents Day 14 wound closure, with larger bubbles indicating lower closure values and smaller bubbles indicating highe closure values.

The day 14 percent closure and the day 14 Histology Resolution Score showed a positive association across all animals tested (Pearson r = 0.68,p = 1.47e^-06^, n = 40, data not shown). Animals with higher surface closure generally tended to have more favorable histology resolution scores. However, animals with similar closure values sometimes differed in histology score, particularly among wounds clustered near the high end of surface closure. This suggests that closure captured an important component of healing, but histology added additional information about tissue-level resolution. Treatment-level means for Day 14 wound closure and Day 14 Histology Resolution Score were z-scored so the two endpoints could be compared on the same relative scale. Values above 0 indicate treatment means above the five-arm cohort mean, while values below 0 indicate treatment means below the overall study mean. The z-scored treatment summary showed strong directional agreement between surface closure and the composite histology score. Group B outperformed Group A animals on both endpoints.

## Conclusions & Discussion

### Specific impact of amnion allograft on wound healing trajectories proprietary and standard processing

The present study demonstrates that differences in tissue processing methodology can influence the biological performance of human amniotic membrane allografts following implantation in a standardized rodent full-thickness excisional wound model. Welch ANOVA across the full five-arm cohort detected a significant overall effect of treatment on the Day 14 Histology Resolution Score, although this pilot was not sized to resolve individual pairwise contrasts under familywise correction. Animals receiving the proprietary processed amniotic membrane exhibited a more favorable wound-healing trajectory than animals treated with a conventionally processed comparator, evident across both gross wound closure and multiple histological endpoints (epidermal, dermal, inflammatory) and reflected in the Day 14 Histology Resolution Score, a composite of microscopic tissue repair endpoints evaluated alongside surface closure. Group B exceeded Group A on both measures: Day 14 Histology Resolution Score 8.94 versus 5.13, and percent wound closure 95.2% versus 89.9% (Table 5).

Epidermal recovery was both more advanced and less injured in Group B. Re-epithelialization was higher (3.75 ± 0.45 vs. 3.44 ± 0.63) and epidermal thickening more pronounced (acanthosis: 3.50 ± 1.03 vs. 3.00 ± 0.63), indicating greater epithelial turnover. Critically, this proliferative activity was achieved with substantially less residual epithelial injury: erosion scores were less than half those of Group A (0.38 ± 0.62 vs. 0.88 ± 0.50), and ulceration was similarly reduced (0.88 ± 1.26 vs. 1.63 ± 1.54), with Group B’s defects limited to scattered superficial involvement rather than extending into the papillary dermis.

The dermal compartment showed a comparable pattern of equivalent regenerative activity alongside reduced tissue injury. Granulation tissue formation was essentially identical between groups (2.94 ± 0.25 in both), and collagen remodeling was closely matched (2.63 ± 0.50 vs. 2.69 ± 0.48), each showing appreciable type I collagen deposition. Angiogenesis (CD31: 3.94 ± 0.25 vs. 4.00 ± 0.00) and epithelial proliferation (Ki-67: 1.69 ± 0.60 vs. 1.50 ± 0.63) were likewise comparable. Against this backdrop of matched regenerative activity, Group B showed markedly less tissue damage: necrosis was less than half that of Group A (0.50 ± 0.73 vs. 1.25 ± 1.13), and hemorrhage was lower as well (0.31 ± 0.48 vs. 0.53 ± 0.63).

The clearest differentiator was acute inflammation, which was more than three-fold lower in Group B (0.50 ± 0.73 vs. 1.63 ± 1.15), with involvement confined to surface scab formation rather than extending into the papillary dermis as seen in Group A. Foreign-body giant cell response was comparably slight in both groups and attributable to incidental rat hair introduction rather than treatment effect. Immunohistochemical findings support this picture of equivalent regenerative signaling achieved with a lower inflammatory burden. CD45 (3.63 ± 0.50 vs. 3.56 ± 0.51) and CD68 (3.19 ± 0.83 in both) were essentially matched, confirming comparable overall leukocytic and macrophage infiltration. CD163 was modestly lower in Group B (3.13 ± 0.62 vs. 3.38 ± 0.62) but remained within a range consistent with active M2-driven matrix remodeling, indicating that Group B reached a comparable regenerative endpoint without relying on as pronounced an M2 macrophage skew.

Taken together, Group B achieved equal or greater epidermal and dermal regenerative activity — matched granulation tissue, collagen remodeling, angiogenesis, and proliferation — while incurring substantially less tissue injury and acute inflammation (necrosis, hemorrhage, erosion, ulceration, and acute inflammation all reduced relative to Group A).. This combination of preserved regenerative capacity with a lower injury and inflammatory burden supports an overall superior day 14 wound healing outcome in Group B. Rather than demonstrating an isolated improvement in a single endpoint, the proprietary allograft produced concordant improvements across multiple independent indicators of tissue repair, including cellular proliferation, tissue remodeling, inflammatory resolution, and overall wound architecture. Although these differences did not reach conventional statistical significance in the present pilot study, the consistency of the observed biological response suggests that the processing methodology may materially influence regenerative performance.

Although the present study was not designed to define molecular mechanisms, these findings are consistent with published observations that amniotic membrane can modulate inflammatory cytokine production, reduce leukocyte activation, and promote transition toward a reparative wound environment [34,35]. Perhaps the most striking observation was improvement in overall tissue organization. Granulation tissue formation was robust and essentially equivalent between groups (2.94 ± 0.25 in both Group A and Group B), and collagen deposition was similarly active in both arms, with comparable alteration scores (2.69 ± 0.48 vs. 2.63 ± 0.50) and appreciable presence of type I collagen fibers indicative of ongoing matrix remodeling. Vascularization, assessed by CD31 immunolabeling, remained high and comparable between groups (4.00 ± 0.00 vs. 3.94 ± 0.25), reflecting sustained angiogenic support of the healing wound bed in both treatment arms. Macrophage-driven remodeling was evident in both groups, with comparable total macrophage infiltration (CD68: 3.19 ± 0.83 in both) and a predominantly M2 (CD163+) phenotype (3.38 ± 0.62 vs. 3.13 ± 0.62), consistent with progression toward a reparative, pro-remodeling wound environment. Taken together, these findings indicate that the proprietary allograft supported equivalent matrix deposition, vascularization, and macrophage-mediated remodeling activity while limiting the excess tissue injury and prolonged acute inflammation observed with standard processing.

### Influence of chemical modification and extracellular matrix preservation

Tissue processing represents an underappreciated determinant of biologic performance. Placental tissue allografts are frequently categorized according to tissue source (amnion, chorion, or amnion/chorion composite), preservation method (cryopreserved, dehydrated, lyophilized), or terminal sterilization strategy [36]. Comparatively less attention has been devoted to understanding how proprietary manufacturing processes alter extracellular matrix composition and ultimately influence biological activity following implantation. Human amniotic membrane functions primarily as a structurally organized extracellular matrix that provides both mechanical support and biological signaling during tissue repair [34,37]. Its regenerative properties depend upon preservation of native collagen architecture, basement membrane integrity, matrix-associated cytokines and growth factors, glycosaminoglycans, adhesive ligands, and endogenous regulators of matrix remodeling [34,36]. Consequently, manufacturing methods that alter these structural or biochemical components may reasonably be expected to influence biological performance following implantation. The present findings support this concept. Although the proprietary processing methodology evaluated here has not yet been mechanistically dissected, the improved wound-healing trajectory observed in vivo suggests that preservation of critical extracellular matrix architecture and associated biological signaling may have contributed to the enhanced regenerative response. Processing methodology influences not merely the rate of wound closure but the quality of regenerated tissue. Restoration of organized extracellular matrix architecture is increasingly recognized as an important endpoint because accelerated wound closure alone does not necessarily predict durable tissue regeneration [38].

One important consideration when interpreting these findings is the potential impact of chemical modification during allograft manufacture. Commercial processing methods may include combinations of dehydration, decellularization, sterilization, and chemical stabilization. Crosslinking chemistries—including glutaraldehyde, carbodiimide (EDC/NHS), genipin, and naturally occurring polyphenols—have been investigated extensively for collagenous biomaterials because they increase mechanical stability and reduce enzymatic degradation [23,39,40,41]. These chemical reactions stabilize collagen by introducing covalent intermolecular crosslinks that increase resistance to proteolysis [42,43]. While improved structural stability may prolong persistence within protease-rich chronic wound environments, excessive crosslinking may also alter several biologically important properties, including extracellular matrix ultrastructure, pore architecture, matrix stiffness, degradation kinetics, cellular attachment, migration, and release of matrix-associated growth factors [41,43]. Numerous biomaterials studies have demonstrated that increasing crosslink density generally decreases cellular infiltration and slows matrix remodeling [41,43]. Likewise, glutaraldehyde crosslinking has long been recognized to increase collagen durability but may also reduce biological remodeling because of residual aldehyde chemistry and increased matrix rigidity [42,27]. Alternative crosslinking approaches such as carbodiimide chemistry or naturally occurring crosslinkers including genipin have been developed partly to improve biocompatibility while maintaining mechanical stability [39,41]. Although the proprietary manufacturing process evaluated here was not specifically designed to investigate individual chemical modifications, the present findings are consistent with the broader concept that preservation of extracellular matrix biology—not simply structural integrity—represents an important determinant of regenerative performance. Future mechanistic studies evaluating collagen ultrastructure, matrix porosity, growth-factor retention, protease susceptibility, and cellular infiltration will be valuable for identifying which processing variables most strongly influence biological activity.

### Clinical implications

The present findings have important implications for interpretation of placental tissue products used in clinical wound care. Placental tissue allografts are often considered interchangeable because they originate from similar donor tissues. However, substantial heterogeneity exists among commercially available products with respect to donor screening, preservation, sterilization, decellularization, dehydration, and proprietary manufacturing methods [36]. Each of these variables has the potential to alter extracellular matrix composition and biological function. The results of this study suggest that placental tissue allografts should not necessarily be regarded as biologically equivalent solely because they share a common tissue source. Rather, manufacturing methodology itself may represent an important determinant of biological performance. If confirmed in larger preclinical studies and prospective clinical investigations, these observations could have implications for evidence-based product selection in wound care and regenerative medicine.

### Study limitations and follow-on study recommendations

Several limitations should be considered when interpreting the present findings. This report presents two arms of a five-arm pilot study, and all inferential statistics were derived from the full five-arm cohort (n = 40). Despite a descriptive trend favoring Group B, the between-group comparison of the histopathology wound resolution score did not reach statistical significance by Games-Howell post hoc testing (p = 0.337), despite a large observed standardized difference between the two arms. At n = 8 per arm, the study was not sized to resolve a pairwise difference of this magnitude under familywise correction.

Using the observed composite effect size as a prospective planning estimate, an unadjusted two-group comparison would require approximately 19 animals per arm to achieve 80% power and approximately 25 per arm to achieve 90% power (α = 0.05, two-tailed). These projections should be treated as a lower bound. The Games-Howell procedure applies a more conservative correction than the unadjusted comparison assumed in this calculation, so a multi-arm design incorporating this correction requires somewhat larger groups; in addition, variance estimates derived from n = 8 are themselves imprecise. Follow-on studies are being sized prospectively from these estimates using pilot variance for the composite score rather than for individual markers.

Histologic scoring in this study was performed by a single observer who was not blinded to treatment group, and no duplicate or adjudicated re-scoring was performed. Unblinded semi-quantitative scoring is a recognized source of observer bias, and this limitation applies to all histology endpoints reported here. Follow-on studies will incorporate standardized, blinded scoring across multiple raters to reduce measurement noise and remove this source of bias.

The current study utilized a healthy rodent excisional wound model rather than models of impaired healing such as diabetic, ischemic, or aged animals [28]. Consequently, any translation to chronic human wounds should be made cautiously and our ongoing studies are focused on chemically and genetically induced diabetic rodent models. Finally, mechanistic molecular analyses—including cytokine profiling, growth factor quantification, matrix metalloproteinase activity, macrophage polarization, transcriptomic analysis, and extracellular matrix proteomics—were beyond the scope of the present investigation but will be incorporated into our current and future follow-on studies.

### Future directions

Future studies should incorporate larger sample sizes, longitudinal molecular analyses, and clinically relevant models of impaired wound healing. Comparative characterization of extracellular matrix composition, collagen ultrastructure, biomechanical properties, protease susceptibility, retained growth-factor profiles, and cellular responses may further elucidate how specific processing methodologies influence regenerative performance. Ultimately, randomized clinical investigations comparing well-characterized placental tissue products will be required to determine whether differences observed preclinically translate into improved patient outcomes.

## Conclusions

In summary, this pilot study demonstrates that manufacturing methodology may substantially influence the biological activity of human amniotic membrane allografts following implantation. Animals treated with the proprietary processed allograft exhibited a consistently more favorable wound-healing trajectory, reflected by improvements in gross wound closure, histological repair, inflammatory resolution, tissue remodeling, and the composite histology resolution index. Although these differences did not achieve statistical significance, the consistency of the observed biological response suggests that preservation of extracellular matrix biology during manufacturing may represent an important determinant of regenerative performance. Collectively, these findings support the concept that placental tissue allografts should not be assumed to be biologically equivalent solely on the basis of tissue source, and they emphasize the importance of rigorous comparative evaluation of manufacturing methodologies when developing and selecting regenerative biomaterials for clinical wound care.

